# Heat-evolved strains of the coral symbiont *Cladocopium proliferum* have diverged in exon usage

**DOI:** 10.64898/2026.11.02.763014

**Authors:** Wenyi Pan, Tianxing Zhou, Thomas W.R. Harrop, Patrick Buerger

## Abstract

Experimental evolution at 31°C produced strains of the coral symbiont *Cladocopium proliferum* (Symbiodiniaceae) that show increased thermal tolerance in culture, and some of them also raise the bleaching tolerance of coral. Here we investigated how exon usage and alternative splicing have diverged in the heat-evolved strains (SS3, SS5, SS8) from their wild-type cultures (WT10) by reanalysing published transcriptomes of the strains in symbiosis with *Acropora kenti* larvae at 27°C using limma, DEXSeq and rMATS. Exon usage separated the strains from an unselected wild-type strain, with 1,308 genes with differential exon usage for SS8, 4,931 for SS5 and 10,464 for SS3, and 59% to 72% of these genes showed no differential expression. SS8, the only strain that also raises the bleaching tolerance of coral larvae, was the closest to the wild type, while a shared set of 757 genes changed in all three heat-evolved strains together. Junction level splicing events (rMATS) did not follow this order (721, 687 and 847 events against WT10), and skipped exons were the most common type, followed by alternative 5’ splice sites. Between SS8 and the non-protective strains, alternative 5’ splice sites were the most common type (91 of 291 events against SS5 and 68 of 207 against SS3), and retained introns were nearly as common as skipped exons. No GO term was enriched in the shared set or the SS8 lists, but both held genes with a possible link to heat tolerance, among them a mitochondrial glutathione peroxidase in the shared set and a glutathione S-transferase in which SS8 differed from the non-protective strains. Future studies of heat-evolved symbionts should therefore include exon level and splicing analyses to resolve the fine scale differences between strains that gene level analyses miss.

## Introduction

Marine heatwaves are the main threat and cause of decline to coral reefs (Hughes et al. 2017), and the Great Barrier Reef had six mass bleaching events between 2016 and 2025 (Ceccarelli et al. 2026). The heat tolerance of a coral depends partly on its microalgal symbionts (Symbiodiniaceae) (Berkelmans and van Oppen 2006; Quigley et al. 2018). When thermal stress exceeds the capacity of the symbiosis, the coral bleaches (loses its symbionts) and likely dies (Glynn 1996).

Assisted evolution may buy time for corals until global warming slows (van Oppen et al. 2015), and one proposed intervention is to evolve the symbionts for heat tolerance in the laboratory (Chakravarti and van Oppen 2018; Nitschke et al. 2024; Scharfenstein et al. 2024). Ten replicate cultures of *Cladocopium proliferum* (formerly *C. goreaui*, type C1^acro^; Butler et al. 2023) from one progenitor culture were selected at 31°C (Chakravarti et al. 2017). After four years, all ten heat-evolved (SS) strains showed enhanced heat tolerance in culture compared to the wild-type (WT). However, only three of the ten strains (SS1, SS7 and SS8) also increased the bleaching tolerance of coral larvae (Buerger et al. 2020), here referred to as protective strains. SS1 and SS8 have since also been shown to enhance the bleaching tolerance of juvenile coral (Quigley and van Oppen 2022; Quigley et al. 2023), and SS8 that of *Acropora spathulata* recruits (Lamb et al. 2026), adult coral (Chan et al. 2023) and the sea anemone *Exaiptasia diaphana* (Chan et al. 2025).

The adaptation of the SS strains to higher temperatures was based to a certain extent on standing genetic variation and recombination (Buerger et al. 2026). Their transcriptomes showed hardly any new mutations and the strains differ in allele frequencies at shared variants (Buerger et al. 2026). However, in terms of molecular mechanisms, it remains mostly unclear what sets the three protective heat-evolved strains (SS1, SS7, and SS8) apart from the unprotective strains. In culture at 27°C the protective strains have, as a group, the lowest intracellular reactive oxygen species and the highest reduced glutathione (Buerger et al. 2023). In coral larvae at 27°C, the protective heat-evolved strain SS8 had carbon fixation genes flagged as enriched in its uniquely upregulated genes compared to the tested unprotective strains (SS3, SS5) (Buerger et al. 2020), and showed three SNPs with significantly different allele frequencies that were associated with carbonic anhydrase genes (Buerger et al. 2026).

Dinoflagellates encode few recognisable transcription factors and rely more on regulation after transcription than other eukaryotes (Bayer et al. 2012; Roy and Morse 2013; Aranda et al. 2016; Zaheri and Morse 2022). This layer can include differential exon usage and alternative splicing. Symbiodiniaceae genes can carry many introns, around 20 per gene, and use GC and GA donor sites as well as the canonical GT (Shoguchi et al. 2013; Roy et al. 2023), and nuclear mRNAs receive a spliced leader by trans splicing (Zhang et al. 2007). The SCF055 assembly of the progenitor culture has 45,322 predicted genes from a workflow modified to recognise these splice sites (Chen et al. 2022). Splicing has been examined in few dinoflagellates, e.g., genome wide in *Amphidinium gibbosum*, where a polyketide synthase gene carried alternatively spliced isoforms (Beedessee et al. 2020), under heat stress in *Prorocentrum cordatum* (Dougan et al. 2023) and in *Durusdinium trenchii* between free living cells and cells in symbiosis with the sea anemone *Exaiptasia diaphana* (as *E. pallida*) at ambient and elevated temperature (Dougan et al. 2024). Exon usage and splicing have not been compared between heat-evolved and wild-type *C. proliferum* strains. Gene level analyses miss it, because they sum the reads of all exons into one count per gene and cannot detect changes in exon usage (Anders et al. 2012).

Here we analyse how far the strains of Buerger et al. (2020) have diverged in exon usage and splicing events (using DEXseq and rMATS), and if any of that divergence supports the protective phenotype of heat-evolved strain SS8. We also identify the genes and exons in which all three heat-evolved strains differ from the wild-type in the same way (shared), and how much of it the gene level analysis misses to inform if future dinoflagellate RNA-seq analyses should consider such a pipeline for fine scale strain differences.

## Methods

### Data and read mapping

We used a published RNA-seq data set (Buerger et al. 2020) consisting of four *C. proliferum* strains in symbiosis with coral larvae (*Acropora kenti*, formerly *Acropora tenuis* larvae; Bridge et al. 2024) at ambient temperatures of 27°C (GEO GSE133082), wild-type (WT10 with 6 samples, WT1 in Buerger et al. 2020), and heat-evolved strains (SS3 with 6 libraries, SS5 (6) and SS8 (5)). In that study, RNA was extracted from a pool of ten larvae and therefore includes coral host and Symbiodiniaceae reads in one sample. Reads were sequenced on a NovaSeq 6000 (2 × 100 bp) with ~87 million reads per sample, out of which mapped 33% to the microalgal symbiont genome, and 53% to the coral host genome (Buerger et al. 2020). In the current study, one SS8 library (40.5, SRR9331955) was left out of the analyses, as in Buerger et al. (2026). It had the fewest read pairs (59.6 million compared to 73 to 101 million for the other libraries) and the highest mismatch rate (0.24% compared to 0.16% to 0.18%; Table S1), it had the lowest share of reads mapped to the two reference genomes combined in Buerger et al. (2020; 61% compared to 86% on average, Database S2), and it was an outlier in our principal component analysis on exon usage. Reads were trimmed with Cutadapt 4.4 (parameters: error rate 0.1, minimum length 50) (Martin 2011) and mapped with STAR 2.7.10b (Dobin et al. 2013) to the soft masked *C. goreaui* SCF055 assembly (GenBank accession GCA_947184155.1, Chen et al. 2022), which was a newer version of the *Cladocopium* reference genome compared to Buerger et al. (2020). Reads were mapped to the genome with the following STAR parameters: sjdbFileChrStartEnd, outSAMmapqUnique 60, outSAMunmapped None, and otherwise default settings. Retained for further analyses (limma voom, DEXseq, and rMATS) were properly paired and uniquely mapped read pairs.

All three analyses used the same five contrasts to measure the divergence of each strain from the wild-type (SS8, SS5 and SS3 each against WT10), and the characteristics of the protective strain from each non-protective strain (SS8 against SS3 and SS8 against SS5). The shared set (SS vs WT) holds the genes with at least one bin that was significant in all three comparisons of SS3, SS5 and SS8 each against WT10 with the same sign of the fold change. For the SS8 comparison against SS3 and SS5 the significant bins were compared if they followed expression in the same direction or opposite directions.

### Gene expression

We used featureCounts 2.1.1 (Liao et al. 2014) to get the gene counts after read mapping. The gene level analysis then repeated the previous pipeline from Buerger et al. (2020) using *limma voom* (Law et al. 2014; Ritchie et al. 2015) and the same filters (filterByExpr, min.count = 40; edgeR, Robinson et al. 2010), TMM normalisation and Benjamini-Hochberg adjusted p-values <0.05 without a fold-change cutoff. The log2 CPM values of the 28,447 genes tested in limma were analysed with a principal component analysis (PCA) on centred, unscaled values in *mixOmics* (Rohart et al. 2017), and with a permutational multivariate analysis of variance on Euclidean distances (PERMANOVA; Anderson 2001; adonis2 in vegan 2.7-1, 999 permutations) followed by a Tukey test on the scores of the first principal component, as in Buerger et al. (2020).

### Exon usage

Exon usage counts per exon bin were generated with dexseq_count.py (DEXSeq 1.56.0, Anders et al. (2012)) on the flattened annotation of dexseq_prepare_annotation.py (789,219 bins, median length 58 bp). Genes with at least 10 read pairs in each of at least 5 libraries (31,470 of the 45,322 annotated genes) entered DEXSeq. 10,239 genes had no uniquely mapped pair in any library. Each gene was fitted with the model ~ sample + exon + strain:exon and a likelihood ratio test for each contrast. Dispersions were estimated under the full model, fold changes with estimateExonFoldChanges, and adjusted p-values with Benjamini-Hochberg. As per our definition, a bin showed differential exon usage (DEU) at an adjusted p-value <0.05 and an absolute log2 fold change of at least 0.5, and a gene was a DEU gene when at least one of its bins was significant for a contrast. Exon usage was analysed with a principal component analysis (PCA) on the log2 CPM of each bin. In each contrast, every DEU gene was classed by the limma result of the same contrast as a) differentially expressed (“DEU+DEG”) if the adjusted p-value for gene expression (DEG, limma) was <0.05, b) exon usage only genes (“DEU”) if the adjusted p-value for DEG was >0.05 (with an adjusted p-value between 0.05 and 0.2 is reported). Genes differentially expressed without differential exon usage are counted as “DEG only”.

### Splicing events

rMATS 4.4.0 (Shen et al. 2014; Wang et al. 2024) ran with --novelSS and --allow-clipping. Events (skipped exon, both exons of a mutually exclusive pair, the extension of the long exon, or the retained intron) were considered significant at FDR <0.05 and an absolute difference in percent spliced in (ΔPSI) above 0.1 on junction counts (JC), and classed as annotated, novel junction (both splice sites annotated, pairing not) or novel splice site from the fromGTF.novelJunction and fromGTF.novelSpliceSite files. Each significant event was related to the DEXSeq bins of the same gene in the same contrast by coordinate overlap.

### Functional summary

GO enrichment was tested with GOseq (Young et al. 2010) as in Buerger et al. (2020) on the respective significant gene lists of this study for each contrast (DEU, DEU+DEG, DEG only, rMATS genes). In addition, listed descriptively were the genes of the two contrasts (SS vs WT, and SS8) with a product name in heat tolerance related families (such as chaperones and heat shock proteins; antioxidant and redox enzymes; photosynthesis and pigments; membrane lipids; osmolytes and ion transport; protein turnover and DNA repair; stress signalling).

## Results

### Gene expression analysis reproduced previous results

Between 22.4% and 42.3% of reads per library mapped uniquely to the symbiont genome (Table 1, S1), as in Buerger et al. (2020, Database S2). The gene level analysis using the published *limma* pipeline produced a similar result, 2.3% of tested genes were differentially expressed between SS8 and SS3 (2020: 3.8%), 6.6% between SS8 and SS5 (6.4%) and 17.7% between SS8 and WT10 (15.2%). The first principal component of gene expression separated WT10 from the three heat-evolved strains (39% of the variance; Figure 1A), strain explained 56.3% of the variation between libraries (PERMANOVA, p = 0.001), and every pair of strains differed on that axis except SS8 and SS5 (Tukey, p = 0.74).

**Table 1.** Libraries and symbiont read depth per strain (22 libraries) showing mean reads (range) per library. SS8 40.5 (SRR9331955) is listed in Table S1 and left out here.

| **Strain** | **Role** | **Samples** | **Read pairs (million)** | **Uniquely mapped pairs (million)** | **Uniquely mapped (%)** | **Pairs on genes (million)** |
|---|---|---|---|---|---|---|
| WT10 | wild-type | 6 | 90.1<br>(80.6 to 97.5) | 25.5<br>(20.9 to 29.6) | 28.4<br>(22.4 to 33.2) | 19.9<br>(16.2 to 23.1) |
| SS3 | heat-evolved,<br>not protective | 6 | 86.3<br>(72.8 to 100.8) | 24.9<br>(20.0 to 30.0) | 28.8<br>(25.2 to 32.3) | 19.4<br>(15.5 to 23.3) |
| SS5 | heat-evolved,<br>not protective | 6 | 86.6<br>(74.6 to 93.9) | 22.0<br>(20.5 to 23.7) | 25.5<br>(22.5 to 29.0) | 17.1<br>(15.8 to 18.3) |
| SS8 | heat-evolved,<br>protective | 4 | 90.1<br>(86.2 to 96.1) | 34.5<br>(31.2 to 36.4) | 38.4<br>(34.7 to 42.3) | 26.9<br>(24.1 to 28.5) |

**Figure 1.**
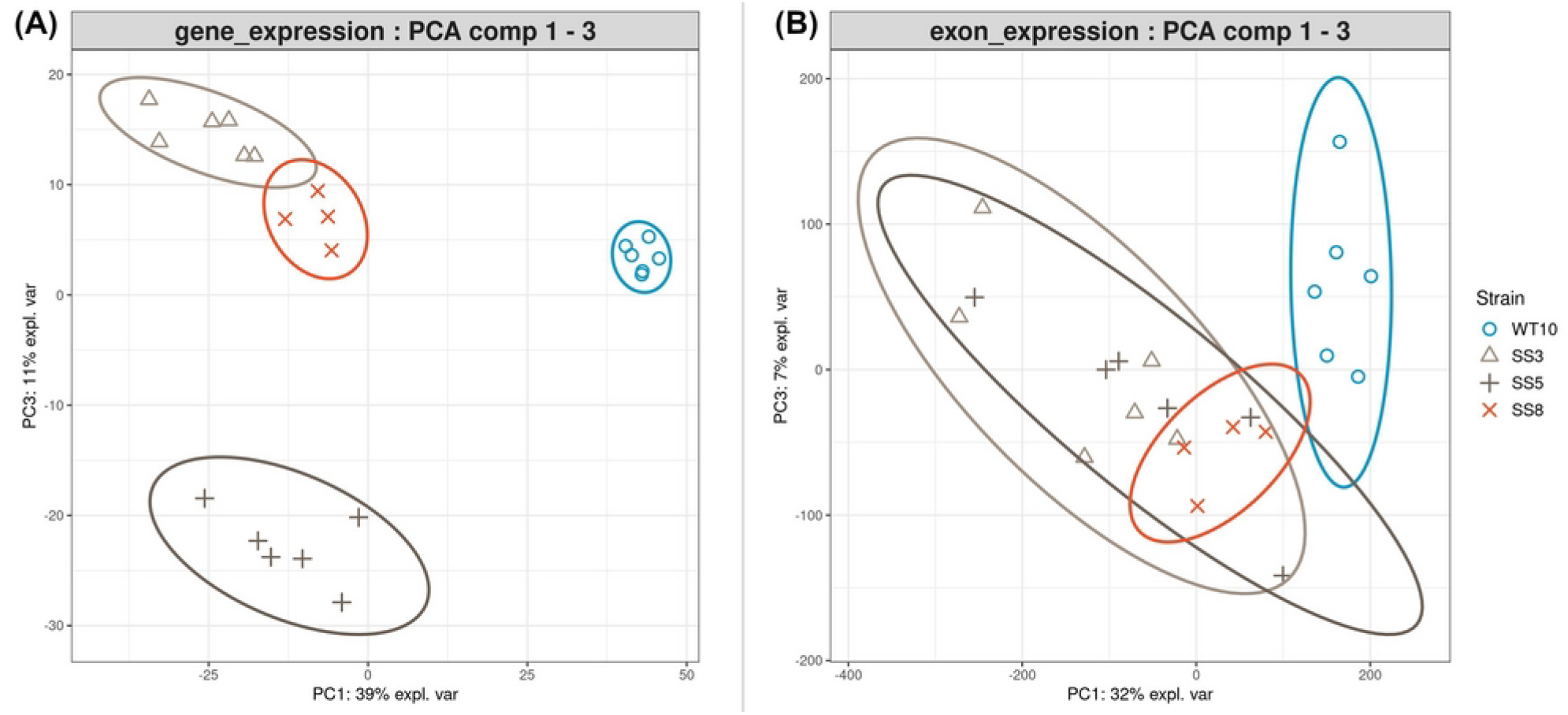
Principal component analysis in *mixOmics* on centred, unscaled values of (A) gene expression, as log2 CPM of the 28,447 genes tested in *limma*, and (B) exon expression, as log2 CPM of the 570,704 DEXSeq bins in the tested genes. Colours and symbols: WT10 blue squares, SS3 light grey circles, SS5 dark grey triangles, SS8 orange diamonds.

**Figure 2.**
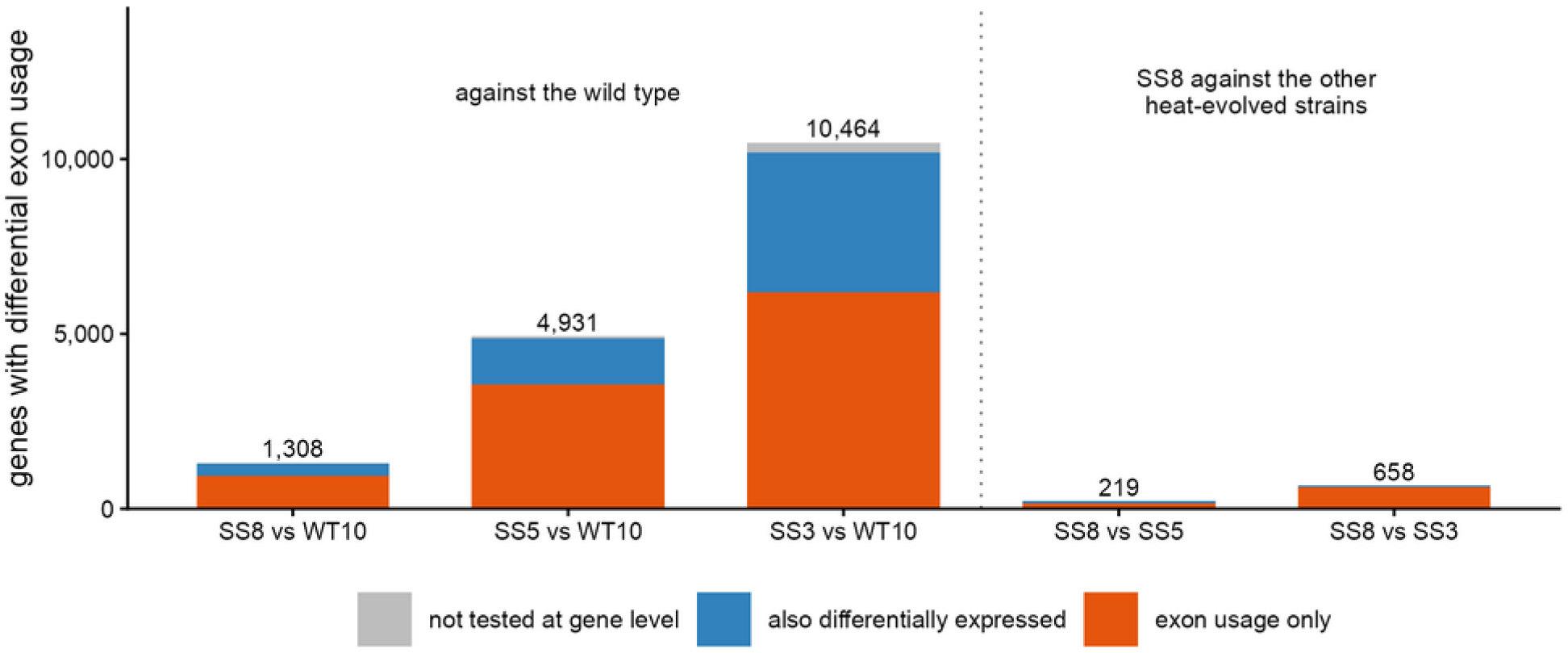
Differential exon usage on the five contrasts. Genes with differential exon usage per DEXSeq contrast, with wild-type against the heat-evolved strains (SS8, SS5 and SS3) and SS8 against each non-protective strain. The categories are determined by the gene level test in the same contrast, also differentially expressed (*limma*, adjusted p <0.05), exon usage only (adjusted p at or above 0.05), or not tested by *limma*.

### Exon usage separates SS from WT

The number of genes with differential exon usage (DEU) against WT10 followed the same order, 1,308 for SS8, 4,931 for SS5 and 10,464 for SS3. DEXSeq tests the reads in each exon bin against the reads in the other exons of the same gene, so a change in overall expression alone does not produce a DEU event (Anders et al. 2012). The two tests therefore measure different things, and their overlap shows how many of the DEU genes a gene level analysis would also have found. Of the 1,308 DEU genes in SS8 against WT10, 939 (72%) were not differentially expressed in the same *limma* contrast (adjusted p ≥ 0.05), as were 3,544 of 4,931 (72%) for SS5 and 6,184 of 10,464 (59%) for SS3. Few of these were near the significance threshold, with only 14% to 27% between 0.05 and 0.2.

The shared set held 757 genes, defined as genes with at least one bin significant in the same direction (higher or lower usage) in each of SS3, SS5 and SS8 against WT10. At gene level, 167 of these were differentially expressed in the same direction in all three strains, and 316 were DEU only. Among the heat-evolved strains, SS8 differed in exon usage from SS3 in 658 genes and from SS5 in 219 genes (824 genes in either contrast, 53 in both). The share of DEU only genes was 93% against SS3 (612 of 658) and 65% against SS5 (143 of 219).

### Splicing events

rMATS found 721 significant events in 526 genes for SS8 against WT10, 687 in 465 genes for SS5 and 847 in 591 genes for SS3, most of them skipped exons followed by alternative 5’ and 3’ splice sites (Figure 3). Most events used a splice site that is not in the gene models (74%, 70% and 73%). Of the events against WT10, 244 were the same event in all three strains with the same direction. SS8 differed in splicing from SS3 in 207 events and from SS5 in 291, with 62 events in both, all in the same direction (higher or lower usage).

**Figure 3.**
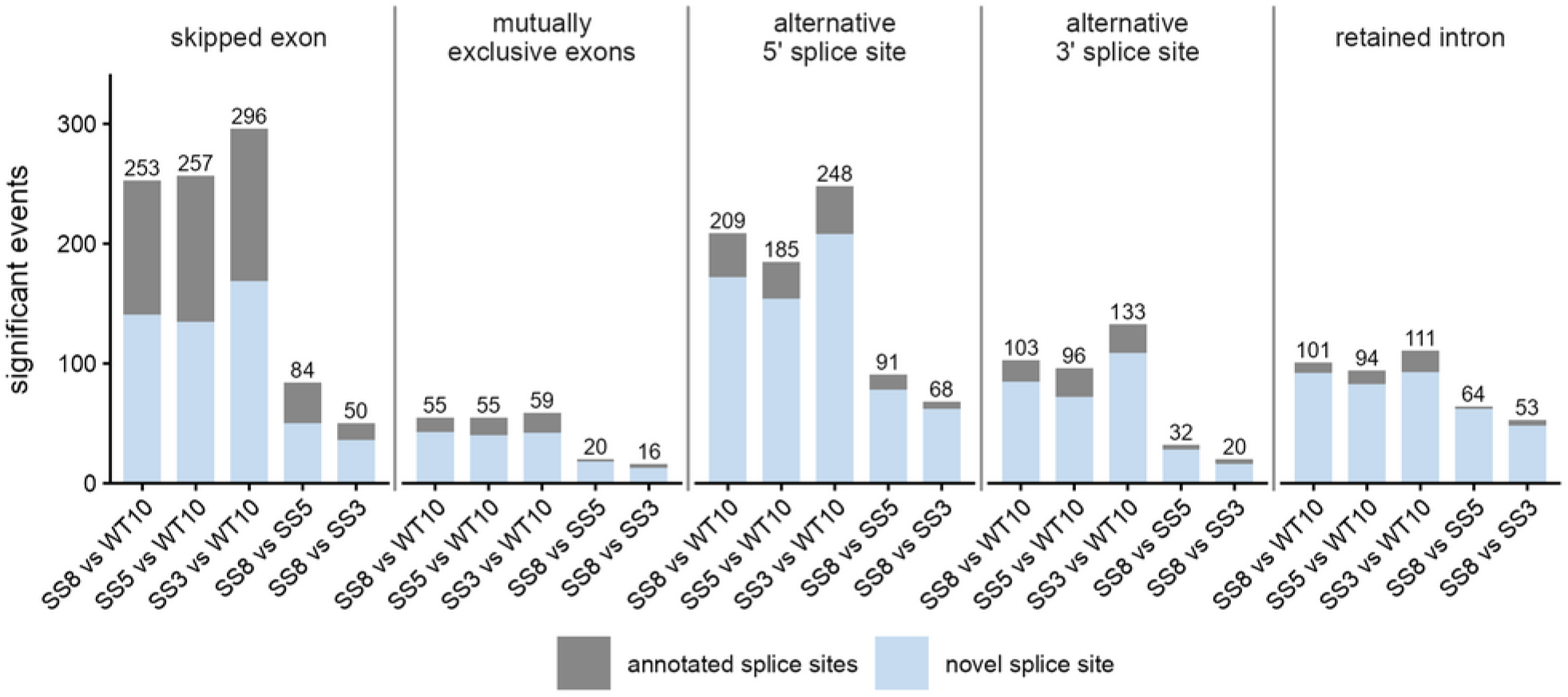
Splicing events from rMATS on the same five contrasts. Significant events (FDR <0.05, absolute ΔPSI above 0.1, junction counts) per type and contrast, split by novel splice sites (blue) and splice sites of the gene models (grey).

LOCUS33148 (transmembrane protein 145) is a DEU only gene with a significant rMATS event at an alternative 3’ splice site. Bin E010 was used in 68% of transcripts in WT10, 26% in SS8, 10% in SS5 and 6% in SS3 (rMATS FDR 6.7 × 10^-18^ for SS8 and below 10^-200^ for SS5 and SS3 against WT10), with no change in total expression (Figure 4A). Example for DEU only with significant rMATS site, exon skipping, LOCUS23058 (hypothetical protein) showed a 15 bp exon included in 64%, 24%, 10% and 14% of transcripts in WT10, SS8, SS5 and SS3 (FDR 1.9 × 10^-59^ for SS8; Figure 4B). LOCUS38398, a zeta-carotene-forming phytoene desaturase, had a lower exon usage of bins E002 and E003 but with higher total expression in all three heat-evolved strains compared to WT (Figure 4C). The fucoxanthin chlorophyll a/c binding protein F gene LOCUS19922 is the opposite case, DEG without DEU or a significant rMATS event, with ~10 fold higher expression in the heat-evolved strains with flat exon usage (Figure 4D).

**Figure 4.**
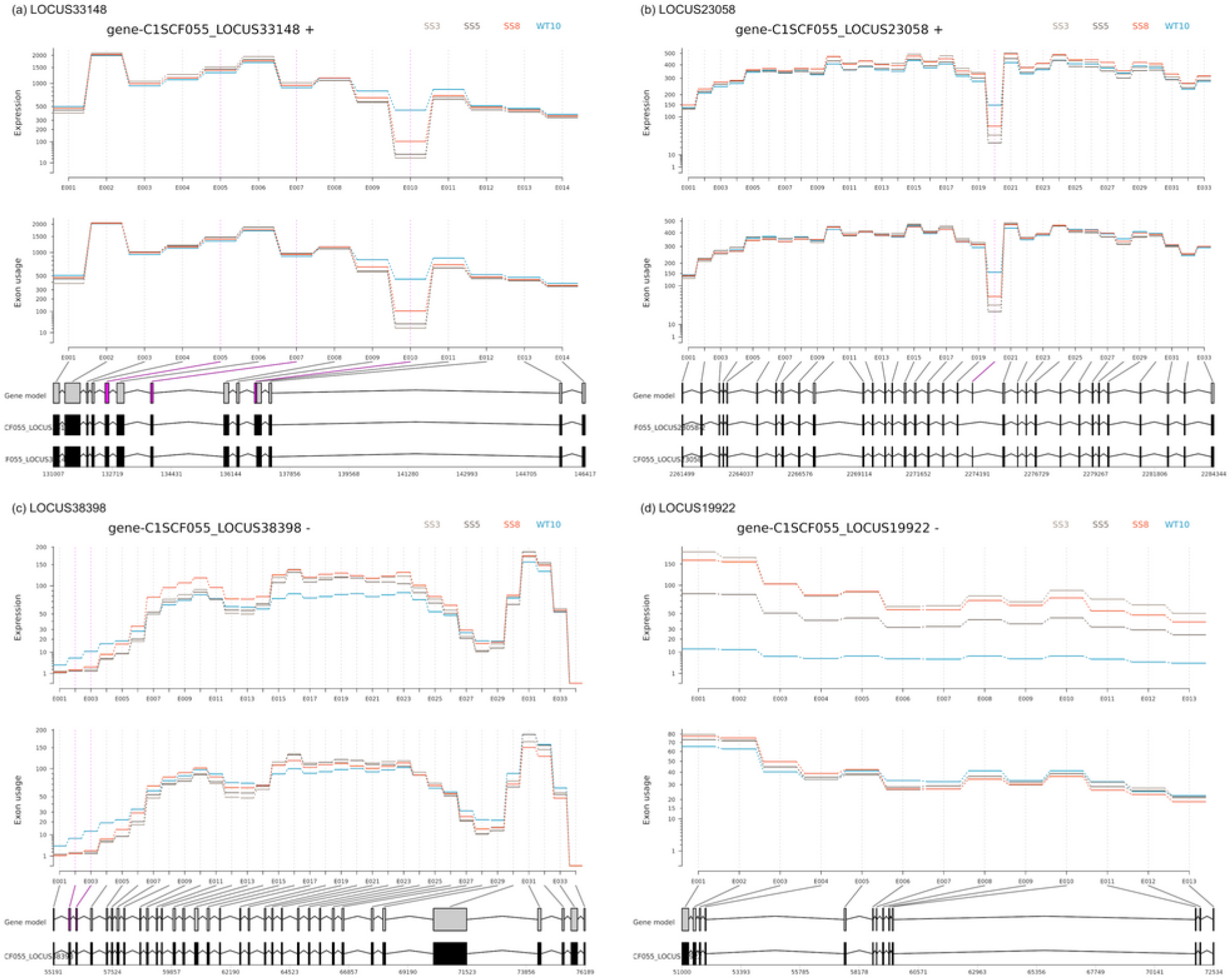
Example genes. plotDEXSeq panels of exon expression (top) and exon usage (bottom) per strain, with the gene model and the bins significant in SS8 against WT10 (pink) below; colours as in Figure 1. (A) LOCUS33148, transmembrane protein 145, alternative 3’ splice site at bin E010. (B) LOCUS23058, hypothetical protein, 15 bp skipped exon at bin E020. (C) LOCUS38398, zeta-carotene-forming phytoene desaturase, changed in both exon usage (bins E002 and E003) and total expression. (D) LOCUS19922, fucoxanthin chlorophyll a/c binding protein F, higher expression in the heat-evolved strains with flat exon usage.

### Heat tolerance related genes follow the same pattern

No GO term was enriched in the shared set, in the SS8 lists or in any other list of this study (GOseq, q <0.05). Of the 757 genes of the shared set, 46 carry a product name in heat tolerance related families, and 15 of the 46 showed no DEG change in any strain. They included for example a mitochondrial glutathione peroxidase (LOCUS30402) for the contrast SS vs WT. Among the genes in which SS8 differs from the non-protective strains, 27 carry such heat stress related gene family, such as a glutathione S-transferase (LOCUS31212; bin E025 1.3 log2 units higher in SS8 than in SS3, Figure 5A). In addition, a subunit of the superoxide-generating NADPH oxidase (LOCUS2272) differed in SS8 from all three other strains (2.9 log2 units lower than WT10, 1.8 higher than SS3, 1.2 higher than SS5, Figure 5B), and a Hsp70 (LOCUS19810) was lower in SS8 than in WT10, SS3 and SS5 (0.67, 0.62 and 0.53 log2 units).

**Figure 5.**
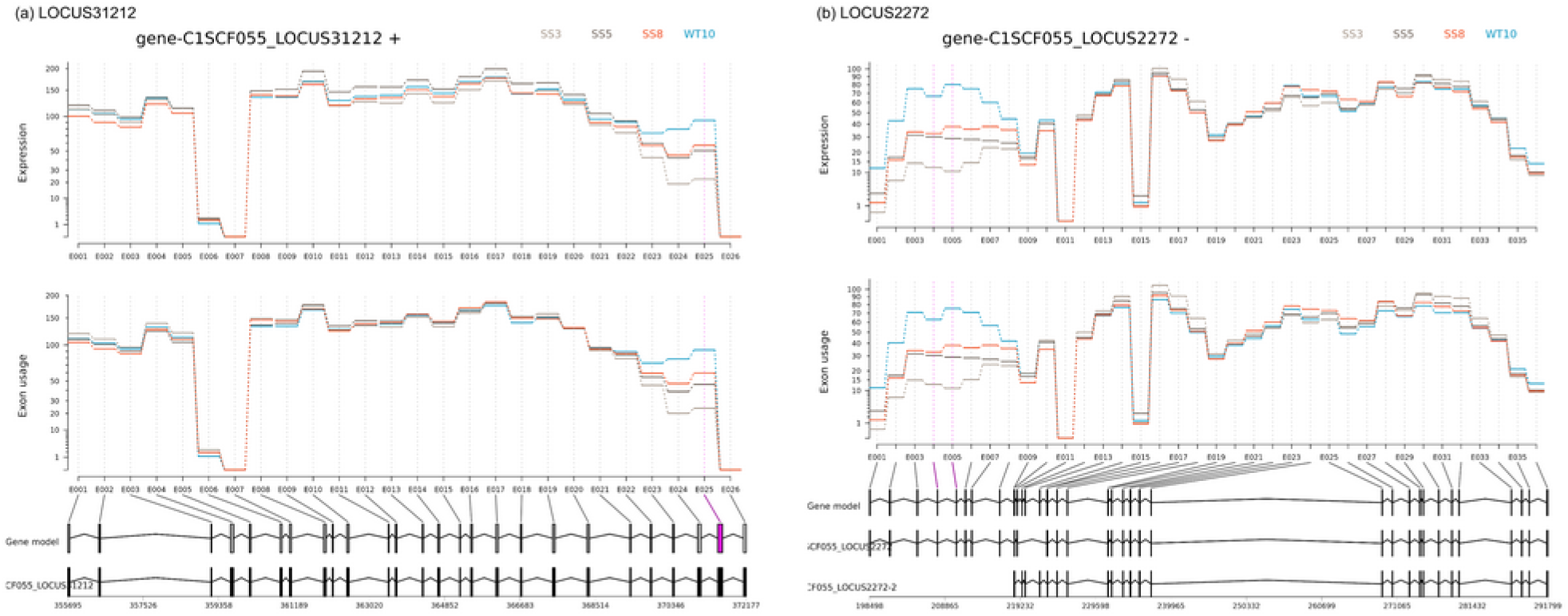
DEXSeq plots of heat tolerance related genes in which SS8 differs in exon usage from the non-protective strains (SS3, SS5). Significant in SS8 against the strains SS3 and SS5 are marked in pink. (a) LOCUS31212, glutathione S-transferase, bin E025, the share of the bin is highest in WT10, then SS8, SS5 and SS3. (b) LOCUS2272, superoxide-generating NADPH oxidase heavy chain subunit, bins E003 to E007 in the same order. SS8 kept the exon usage of the wild-type more closely than SS5 and SS3, and none of the two genes changed in total expression in SS8 against WT10.

## Discussion

Heat-evolved strains of *C. proliferum* have been tested in coral larvae, juvenile corals, adult corals and sea anemones (Buerger et al. 2020; Quigley and van Oppen 2022; Quigley et al. 2023; Lamb et al. 2026; Chan et al. 2023, 2025). Their transcriptomes in symbiosis had so far been analysed at gene level only (Buerger et al. 2020), and exon level analysis of the same data shows that this missed most of the divergence. Three heat-evolved strains differed in exon usage in 1,308 to 10,464 genes from a wild-type strain kept at 27°C for the same four years, and 59% to 72% of these genes showed no difference in overall expression. The strains formed a gradient of divergence from the wild type, with SS8 the nearest and SS3 the furthest, and 757 genes changed in the same direction in all three heat-evolved strains.

The three analyses do not agree on which non-protective heat-evolved strain is closer to SS8. At gene level, SS8 differed from SS5 in more genes than from SS3 (6.6% against 2.3% of tested genes), however, SS8 and SS5 did not differ on the first principal component. At exon level SS8 differed from SS3 in three times as many genes as from SS5 (658 compared to 219), while splicing events were reverse again (207 against SS3, 291 against SS5). The strain that looks closest to SS8 therefore depends on the measured parameter, which is a further sign that gene level and exon level analyses capture different layers with their own resolution.

Divergence at this level is what the biology of dinoflagellates predicts, given how little of their regulation runs through transcription factors (Roy and Morse 2013; Zaheri and Morse 2022). Many of their genes change little in expression under heat, and expression differences between strains are often fixed instead of induced (Barshis et al. 2014; Buerger et al. 2020). In *Prorocentrum cordatum*, 1,820 of 85,849 genes differed in exon usage in an experiment at 20, 26 and 30°C (Dougan et al. 2023). In *Durusdinium trenchii*, among duplicated gene pairs with opposite expression in free living and symbiotic cells, the copy specialised for symbiosis had fewer exons and less differential exon usage than its free living counterpart, which the authors interpreted as streamlining for the symbiotic lifestyle (Dougan et al. 2024). In *Amphidinium gibbosum*, one exon of a polyketide synthase gene changed in usage under nitrogen starvation (Beedessee et al. 2020). In *Symbiodinium microadriaticum*, a subset of genes showed condition specific RNA editing under cold, heat and dark stress (Liew et al. 2017). In vertebrates, splicing diverges faster than gene expression between lineages (Barbosa-Morais et al. 2012; Merkin et al. 2012), and most species-specific splicing patterns are directed by sequences in the gene itself (Barbosa-Morais et al. 2012). At the same time, part of what RNA-seq detects can be splicing noise, and many alternative transcripts may never be translated (Pickrell et al. 2010; Tress et al. 2017). Nevertheless, some of the genes here with differential exon usage and with relevance to heat tolerance related gene families might be candidates for early steps of adaptation to 31°C.

Against WT10, skipped exons were the most common event type, followed by alternative 5’ and 3’ splice sites. Skipped exons also dominated a genome wide survey of alternative splicing in *Amphidinium gibbosum*, to our knowledge the only other dinoflagellate analysed with rMATS (6,970 events in 5,417 genes; 77.2% skipped exons, 11.3% alternative 5’ and 6.8% alternative 3’ splice sites; Beedessee et al. 2020). Between SS8 and the non-protective strains, alternative 5’ splice sites were the most common type (91 of 291 events against SS5 and 68 of 207 against SS3), and retained introns were nearly as common as skipped exons. In *A. gibbosum*, retained introns and mutually exclusive exons together made up the remaining 4.7% of events (Beedessee et al. 2020). In mammals, retained introns can reflect slower splicing, or transcripts held in the nucleus or targeted for decay (Braunschweig et al. 2014; Boutz et al. 2015). Our data cannot separate these, or show whether the same applies in dinoflagellates.

Most events against WT10 (70% to 74%) used a splice site absent from the gene models. The isoforms in the SCF055 models were added from transcript evidence, including long reads from cultured cells in exponential growth (55,505 transcripts for 45,322 genes; Chen et al. 2022). Splice sites used only in symbiosis, or in transcripts too rare to be captured, would be missing from them. A novel site used differently between strains is likely still a strain difference, because all strains were mapped to the same reference.

The three heat-evolved strains changed 757 genes in the same direction. Selection at 31°C may have acted in parallel on the same standing variants, because the strains came from one progenitor culture, carried few new mutations and differed in allele frequencies at shared variants (Buerger et al. 2026). Some of these changes may instead have hitchhiked with selected alleles, and some may reflect drift in WT10. The overlap between these exons (DEU, and splicing events) and the variants that shifted during selection would clarify some of their impact.

Four years of selection at 31°C left three closely related strains of *C. proliferum* with different exon usage and splicing patterns, and most of that difference is not visible at gene level. Earlier gene expression analyses of heat-evolved strains from this experiment summed the reads of each gene (Chakravarti et al. 2020; Buerger et al. 2020) and could not detect differences within a gene. Since exon level analysis needs no new sequencing, it should be part of how evolved symbionts are characterised. Long read transcriptomes of these strains under heat stress, and the overlap with the variants that shifted during selection, would show if exon usage matters for heat tolerance.

## Author statements

### Author contributions

W.P.: formal analysis (exon usage and splicing, first analysis), writing (review and editing). T.Z.: formal analysis (read mapping), writing (review and editing). P.B. and T.H.: conceptualisation, formal analysis (final analysis), writing (review and editing), supervision, writing (original draft).

## Acknowledgements

We thank Cheong Xin Chan for the SCF055 functional annotation. Analyses ran on the National Computational Infrastructure (NCI Gadi, project rm18 and d85) and on Galaxy Australia.

