## Supplementary Materials for "Heat-evolved strains of the coral symbiont *Cladocopium proliferum* have diverged in exon usage"

### Supplementary Tables

**Table S1.** Overview and mapping of the 23 RNA-seq libraries. Read pairs from the SRA record; mapping from STAR; pairs on genes from featureCounts (reverse strand). SRR9331955 (SS8 40.5, marked) is left out of the main analysis.

| Run | GEO title | Strain | Read pairs (million) | Uniquely mapped (%) | Multimapped (%) | Unmapped, too short (%) | Pairs on genes (million) | Mismatch rate (%) | Non canonical junction reads (%) |
|---|---|---|---|---|---|---|---|---|---|
| SRR9331960 | WT1 27_50.1 | WT10 | 80.6 | 28.32 | 4.81 | 65.68 | 17.7 | 0.18 | 16.91 |
| SRR9331961 | WT1 27_50.5 | WT10 | 89.2 | 33.24 | 5.55 | 59.84 | 23.1 | 0.17 | 16.94 |
| SRR9331962 | WT1 27_50.6 | WT10 | 90.2 | 25.18 | 3.94 | 69.96 | 17.7 | 0.17 | 16.92 |
| SRR9331963 | WT1 27_50.7 | WT10 | 89.8 | 31.60 | 5.37 | 61.68 | 22.1 | 0.17 | 16.92 |
| SRR9331964 | WT1 27_50.8 | WT10 | 97.5 | 29.44 | 4.61 | 64.87 | 22.4 | 0.17 | 16.83 |
| SRR9331965 | WT1 27_50.9 | WT10 | 93.5 | 22.37 | 3.80 | 72.90 | 16.2 | 0.16 | 17.02 |
| SRR9331943 | SS3 27_14.3 | SS3 | 72.8 | 27.48 | 4.29 | 67.24 | 15.5 | 0.17 | 17.12 |
| SRR9331944 | SS3 27_14.4 | SS3 | 90.9 | 25.25 | 4.30 | 69.37 | 17.8 | 0.18 | 17.15 |
| SRR9331945 | SS3 27_14.5 | SS3 | 85.4 | 28.96 | 4.47 | 65.52 | 19.3 | 0.17 | 16.93 |
| SRR9331946 | SS3 27_14.6 | SS3 | 84.1 | 29.26 | 5.10 | 64.33 | 19.2 | 0.17 | 16.95 |
| SRR9331947 | SS3 27_14.7 | SS3 | 83.6 | 32.32 | 5.41 | 60.91 | 21.0 | 0.17 | 16.92 |
| SRR9331948 | SS3 27_14.8 | SS3 | 100.8 | 29.77 | 4.85 | 64.20 | 23.3 | 0.18 | 17.04 |
| SRR9331949 | SS5 27_25.1 | SS5 | 81.5 | 27.27 | 4.25 | 67.48 | 17.4 | 0.17 | 16.83 |
| SRR9331950 | SS5 27_25.5 | SS5 | 74.6 | 28.97 | 5.03 | 64.72 | 16.6 | 0.18 | 17.04 |
| SRR9331951 | SS5 27_25.6 | SS5 | 87.2 | 23.48 | 3.74 | 71.90 | 15.8 | 0.18 | 17.12 |
| SRR9331952 | SS5 27_25.7 | SS5 | 90.0 | 26.28 | 4.48 | 68.13 | 18.3 | 0.17 | 17.07 |
| SRR9331953 | SS5 27_25.8 | SS5 | 93.9 | 24.80 | 4.18 | 69.98 | 18.1 | 0.17 | 17.01 |
| SRR9331954 | SS5 27_25.9 | SS5 | 92.1 | 22.47 | 3.89 | 72.65 | 16.0 | 0.18 | 16.92 |
| SRR9331955 | SS8 27_40.5 | SS8 (left out) | 59.6 | 32.22 | 5.40 | 60.98 | 15.0 | 0.24 | 16.70 |
| SRR9331956 | SS8 27_40.6 | SS8 | 89.9 | 34.67 | 6.05 | 57.72 | 24.1 | 0.17 | 16.95 |
| SRR9331957 | SS8 27_40.7 | SS8 | 86.2 | 42.27 | 6.72 | 49.36 | 28.5 | 0.17 | 16.87 |

| Run | GEO title | Strain | Read pairs (million) | Uniquely mapped (%) | Multimapped (%) | Unmapped, too short (%) | Pairs on genes (million) | Mismatch rate (%) | Non canonical junction reads (%) |
|---|---|---|---|---|---|---|---|---|---|
| SRR9331958 | SS8 27_40.8 | SS8 | 96.1 | 36.42 | 6.22 | 55.76 | 27.2 | 0.17 | 16.92 |
| SRR9331959 | SS8 27_40.9 | SS8 | 88.3 | 40.24 | 6.91 | 51.07 | 27.6 | 0.17 | 16.96 |
